# Intact learning and memory in mice incapable of de novo myelination

**DOI:** 10.64898/2026.06.12.730403

**Authors:** Matthew Swire, Stuart G Nayar, Yi Jiang, Adam Halko, Marcus Lloyd, Katsutoshi Ogasawara, Koujiro Tohyama, Thomas Philips, Jeffrey D Rothstein, Huiliang Li, William D Richardson

**Affiliations:** Wolfson Institute for Biomedical Research, University College London, Gower Street, London WC1E 6BT, UK; Department of Physiology, Iwate Medical University, 1-1-1 Idaidori, Yahaba-cho, Shiwa-gun, Iwate 028-3694, Japan; Johns Hopkins University School of Medicine, 855 North Wolfe Street, Baltimore, MD 21205, USA

**Author notes:** Correspondence to:* Prof William D Richardson or Prof Huiliang Li. Wolfson Sensory, Pain and Regeneration Centre (SPaRC), King’s College London, Hodgkin Building, Guy’s Campus, London SE1 1UL, UK. Oxford PharmaGenesis, Tubney, Oxford OX13 5QJ, UK. Institute of Ophthalmology, University College London, London, UK. QurAlis, 35 Cambridgepark Drive, Cambridge, MA 02140, USA. these authors made equivalent contributions (Matthew Swire, Stuart G Nayar). joint senior authors (Huiliang Li, William D Richardson).

**Keywords:** myelin basic protein, MBP, conditional knockout, Shiverer mutant mouse, node of Ranvier, CASPR, NaV, motor learning, fear conditioning, memory consolidation

## Abstract

Motor skill learning stimulates and requires generation of oligodendrocytes (OLs) from their precursors (OLPs) in the adult mouse brain, but the functional role(s) of the newly formed OLs is not known. We asked whether new compact myelin sheaths are required, by genetic block of myelin basic protein (MBP) synthesis in adult OLPs and their newly-differentiating OL progeny, using tamoxifen-inducible Cre-*lox* recombination. Newly-differentiating OLs in these *Mbp-cKO* mice are unable to assemble compact myelin or normal nodes of Ranvier. Despite this, *Mbp-cKOs* learned a motor skill just as well as their wild type littermates. They also demonstrated normal contextual fear conditioning. Therefore, neither motor nor fear learning depends on rapid saltatory conduction in newly-myelinated circuits. *Mbp-cKOs* also formed normal long-term motor and contextual fear memories. Moreover, OL lineage-specific knockout of *Monocarboxylate transporter-1 (Mct1),* believed to be responsible for transferring metabolic substrates from OLs into axons, did not affect learning or memory consolidation. *Myelin regulatory factor (Myrf)-cKOs*, in which newly-differentiating OLs die and are rapidly eliminated, confirmed that newly formed OLs are required for learning and memory. Together, the data suggest that learning and memory depends on a non-canonical property of pre-myelinating or myelinating OLs, distinct from myelin’s cardinal role in speeding action potentials.

## Introduction

The oligodendrocyte (OL) lineage exhibits marked plasticity in healthy adult mice, particularly during learning experiences such as motor skill learning,^1–6^ fear and spatial memory consolidation, ^7–9^ and T-maze and radial maze learning tasks that rely on working memory.^10^ During training, a subset of adult OL precursors (OLPs), most of which are normally resting in the G1-phase of the cell cycle, are triggered to exit the division cycle and differentiate directly into OLs, many of which go on to form new myelin sheaths. The generation of new OLs leads to a transient depletion of the OLP population and the surviving OLPs mount a robust homeostatic response, ^11–14^ moving rapidly from G1- to S-phase and undergoing cell division to repopulate the precursor pool. This dynamic cell behaviour can be dramatic; we found that, in the anterior corpus callosum, the proportion of OLPs that enters S-phase in a mouse undergoing radial maze training is directly proportional to individual performance and can exceed 90% of the OLP population in the best-performing animals.^10^

When we blocked OLP differentiation in adult mice by conditionally ablating the *Myelin regulatory factor (Myrf)* gene in OLPs using Cre-*lox* recombination (with *Pdgfra-CreER^T^*^2^), this impaired their ability to learn a new motor skill (running on a “complex wheel” with unequally spaced rungs),^1^ or to improve their performance in maze tasks that rely on working memory.^10^ Other labs used *Ng2-CreER^TM^* or *Pdgfra-CreER^T2^* for conditional knockout of *Myrf* and found that the mice were impaired in remote (30 day) recall of fear memory or spatial memory, although learning per se was not affected.^7–9^ These findings have contributed to a working hypothesis that generation of new myelinating OLs is required for some forms of learning as well as for consolidation of long-term memories.^10^

*Myrf* encodes a transcription factor that is absolutely required for successful OL differentiation. In its absence, OLPs embark on an abortive differentiation program, die as pre-myelinating OLs (pmOLs) and are rapidly cleared away by phagocytosis.^1,15^ Consequently, almost no newly-differentiating OLs can be detected in *Pdgfra-CreER^T2^: Myrf ^(fl/fl)^* (*Pdgfra-Myrf-cKO*) mice compared to control littermates^1,2^ It follows that the behavioural phenotype of *Pdgfra-Myrf-cKOs* may not be attributed solely to loss of new myelin sheath production along with their canonical role of accelerating action potentials; learning deficits caused by *Myrf* deletion could equally result from loss of some non-canonical function of new myelinating or pre-myelinating OLs.

Here, we aimed to determine whether newly-generated, compacted myelin sheaths are important for learning and consolidation of novel motor skills and contextual fear cues. We used mouse genetics to interfere with the biology and function of newly-formed OLs without causing their early death and removal. We generated new *Mbp^(fl/fl)^* mice that were unable to produce *Myelin basic protein (Mbp)* mRNA or MBP protein – hence normal compact myelin – following Cre-recombination. When *Mbp^(fl/fl)^* mice were crossed with *Pdgfra-CreER^T2^, Ng2-CreER^TM^* or *Sox10-iCreER^T2^*, myelin developed normally in the double-transgenic *Mbp^(fl/fl)^* offspring until tamoxifen was administered in young adulthood, after which differentiated OLs continued to be generated but were unable to synthesize compact myelin sheaths with normal nodes of Ranvier. These “new-myelin-deficient” mice were able to learn to run on the complex wheel just as fluently as normal littermate controls and retained this skill over at least a month, leading us to conclude that the canonical function of the newly-formed OLs – to enable rapid saltatory conduction along newly-myelinated axonal segments – is not essential for motor skill learning or memory. The mice also displayed normal fear-conditioning responses and long-term fear memory. These findings suggest that a non-canonical function of newly-formed OLs is required to support learning and memory.

## Results

### Developmental block of myelin basic protein transcription mimics “shiverer” mutation

The *Mbp* gene, encoding myelin basic protein (MBP), a major structural protein of myelin, is transcribed as part of a larger transcription unit *Golli-Mbp* comprising 11 exons within a >100kb region of mouse chromosome 18 ^16^. We used gene targeting to insert *loxP* sites into intron 4 and exon 5c, flanking the *Mbp* transcription start site in exon 5b, so as to eliminate *Mbp-*specific transcripts by Cre-*lox* recombination without affecting the longer *Golli* transcripts that initiate upstream of exon 1 (**Fig. 1A**). This approach is similar to that described by Meschkat et al.,^17^ who inserted *loxP* sites into intron 4 and intron 5 of *Golli-Mbp* flanking the *Mbp* promoter (**Fig. 1A**). Therefore, our *Mbp^fl^* allele and that of Meschkat et al.^17^, though subtly different in design, are expected to have similar properties.

**Figure 1.**
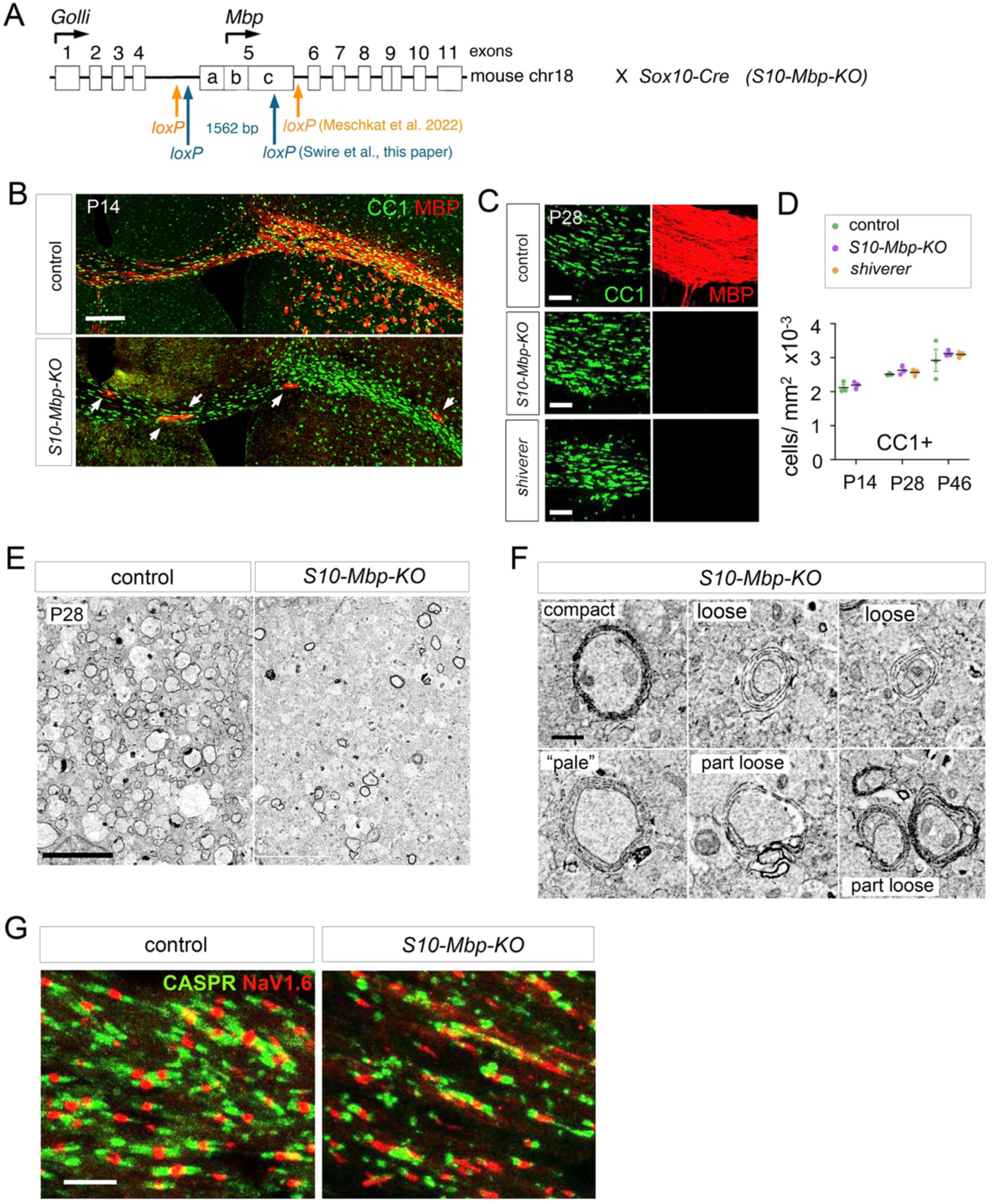
Efficient block of MBP synthesis in constitutive *S10-Mbp-KO* mice and comparison with *shiverer* mutant mice. (**A**) Structure of the *Golli-Mbp* gene showing the internal *Mbp* transcription start site and the insertion sites of *loxP* recognition sites for Cre recombinase in our *Mbp^fl^* mice, used in the present study, and in the *Mbp^fl^* mice made and described by Meschkat et al. (2022). Our *Mbp^(fl/fl)^* mice were crossed with female *Sox10-Cre: Mbp^(fl/+)^* to generate constitutive *S10-Mbp-KOs* and *Mbp^(fl/+)^* littermates. (**B**) Immunolabelling of P14 mouse coronal forebrain sections at the level of the motor cortex, containing subcortical white matter (SCWM) from the midline of the brain (left) to one lateral extremity (right), with motor cortex above and striatum below (dorsal up). In controls, most CC1^+^ OLs (green) are also MBP^+^ (red) in SCWM and striatum (upper panel). In *S10-Mbp-KOs,* most CC1^+^ OLs lack MBP immunolabelling, except for very occasional cells (<1%) that escaped recombination (white arrows). (**C**) Enlarged images of the corpus callosum, with an equivalent region of *Mbp^(shi/shi)^* brain for comparison. The density of CC1^+^ differentiated OLs appears similar in all three specimens. (**D**) Cell counts confirm that there are similar number-densities of CC1^+^ OLs in *S10-Mbp-KO, Mbp^(fl/+)^*, *Mbp^(shi/shi)^* and control SCWM at P14-P46. (**E**) Wide-field backscatter EM of sagittal sections of SCWM show that *S10-Mbp-KO* mice have far fewer cross sections of normal electron-dense myelin sheaths compared to their control littermates. (**F**) Higher-magnification examples of “loose” or “pale” uncompacted myelin sheaths in *S10-Mbp-KO.* (**G**) Double-immunolabelling for CASPR (green) and NaV1.6 (red) reveals nodes of Ranvier with typical NaV clusters flanked symmetrically by CASPR^+^ paranodes in control SCWM, but disorganized nodal structures, frequently with abnormally short paranodes, in *S10-Mbp-KO* SWCM. *Scale bars:* (**B**) 1 mm, (**C**) 200 µm, (**E**) 5 µm, (**F**) 2 µm, (**G**) 5 µm.

When we crossed male *Mbp^(fl/fl)^* mice with female *Sox10-Cre: Mbp^(fl/+)^* BAC transgenics, which express Cre constitutively in all developmental stages of the OL lineage, their double transgenic *Sox10-Cre: Mbp^(fl/fl)^* (*S10-Mbp-KO*) offspring developed normal numbers of differentiated OLs that expressed adematous polyposis coli (APC), recognized by monoclonal CC1 (**Fig. 1B-D**).

However, the vast majority of CC1^+^ OLs did not express MBP protein (**Fig. 1B, C**) and formed many fewer compact myelin sheaths than normal (**Fig. 1E, F**). Frequent examples of loosely-ensheathed axons were observed in electron microscope images (**Fig. 1F**), similar to the spontaneous *Mbp* deletion mutant *shiverer* (*Mbp^(shi/shi)^*).^18^ In addition, the white and grey matter of *S10-Mbp-KO* mice contained abnormal, disorganized nodes of Ranvier (**Fig. 1G**). Although some NaV1.6 sodium channel clusters still formed in *S10-Mbp-KOs*, they were more variable in size than in control *Sox10-Cre: Mbp^(fl/+)^*littermates and they were usually associated with abnormally short CASPR1^+^ paranodes (**Fig. 1G**), as described before for *Mbp^(shi/shi)^.*^19,20^ Constitutive *S10-Mbp-KO* mice developed apparently normally until the second postnatal week, when they developed intentional tremor resembling that described for *Mbp^(shi/shi)^*(**supplementary Movies S1, S2**).^21^ We concluded that our *Mbp^fl^* allele can be recombined in an efficient and targeted manner, effectively preventing the synthesis of MBP protein and compact myelin in OL lineage cells, and is therefore a potentially powerful tool for investigating the role of myelin in physiological processes including learning and memory formation.

*Preventing production of Mbp^+^ myelinating OLs during adulthood does not inhibit skill learning* To test the idea that generation of new compact myelin is required for motor skill learning during adulthood, we crossed *Mbp^(fl/fl)^* with *Ng2-CreER^TM^* mice^22^ to generate conditional knockout (cKO) offspring. We administered tamoxifen to young adult (P60–P63) *Ng2-CreER^TM^: Mbp^(fl/fl)^* (*Ng2-Mbp-cKO*) mice and littermate controls (*Ng2-CreER^TM^: Mbp^(fl/+)^*), provided EdU in their drinking water for 10 days P80–P90 (**Fig. 2A**), then visualized newly-generated *Mbp*^+^ EdU^+^ OLs in the corpus callosum (CC) by fluorescence in situ hybridization (FISH) for *Mbp* combined with “click-it” histochemistry for EdU (**Fig. 2B**). The total number-density of EdU^+^ cells was not significantly altered in *Ng2-Mbp-cKO* mice relative to littermate controls, but the density of *Mbp^+^*, EdU^+^ newly-generated OLs was reduced by ∼70% (118 ± 4 cells/ mm^2^ in *Mbp^(fl/+)^* vs 36 ± 4 cells/mm^2^ in *Mbp^(fl/fl)^*, n = 3 mice of each genotype), indicating that both *Mbp^fl^* alleles had been deleted in ∼70% of OLPs (**Fig. 2B, C**). These *Mbp-*negative OLPs should differentiate into OLs that can survive long-term but are incapable of synthesizing MBP protein, hence unable to generate compact myelin.

**Figure 2.**
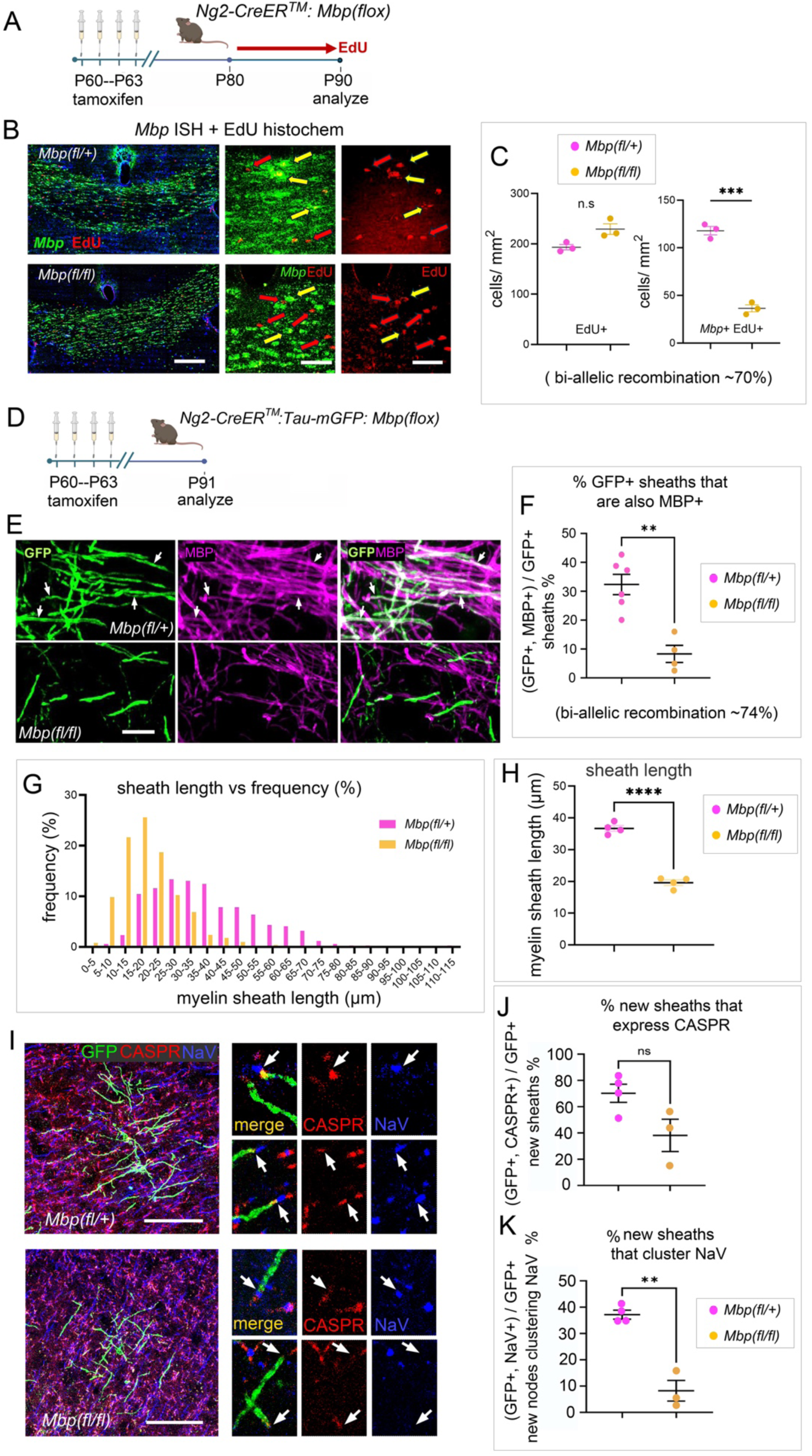
Conditional block of *Mbp* transcription in adult OPCs using *Ng2-CreER^TM^* strongly inhibits production of MBP^+^ OLs and normal myelinated axon units. (**A**) Tamoxifen was administered to to *Ng2-CreER^TM^: Mbp^(fl/fl)^*mice (*Ng2-Mbp-cKOs*) and their *Mbp^(fl/+)^* littermates from P60–P63 and EdU was given in the drinking water for 10 days P80–P90 to allow detection of dividing OPCs and their differentiated OL progeny. (**B**) Sections of *Ng2-Mbp-cKO* and control brains showing the corpus callosum beneath the motor cortex, labelled for *Mbp* RNA by FISH and for EdU by histochemistry. Separate fluorescence channels for *Mbp* and EdU are shown on the right, at higher magnification. In control sections, many *Mbp^+^* OL cell bodies (green) co-labelled for EdU (red), showing that they had differentiated from OPCs before tamoxifen was given (*yellow arrows*), while other EdU^+^ cell nuclei were *Mbp-*negative, indicating that these were OLPs that had not yet differentiated and initiated *Mbp* expression (*red arrows).* In *Ng2-Mbp-cKO* mice there were fewer double-positive OLs, indicating that *Mbp* had been deleted in a proportion of OLPs prior to differentiation (*red arrows*). (**C**) The total number of EdU^+^ OL lineage cells was not significantly changed in *Ng2-Mbp-cKO* mice, but the number of GFP^+^, *Mbp^+^* double-positive cells decreased compared with littermate controls, indicating that the efficiency of *Mbp* deletion in OLPs was ∼70%. (**D, E**) For another estimate of recombination efficiency, we administered tamoxifen to *Ng2-CreER^TM^: Mbp^(fl/fl)^: Tau-mGFP* mice at P60–P63, immunolabelled for GFP (green) and MBP (magenta) and counted double-labelled new sheaths (GFP^+^, MBP^+^) and single-labelled GFP^+^ new sheaths 1 month later, on P94. (**F**) The fraction of new GFP^+^ sheaths in the motor cortex that co-labelled for MBP (white) was lower in *Mbp^(fl/fl)^* mice than in their *Mbp^(fl/+)^* littermates, indicating that recombination efficiency was around 74%. (**G**, **H**) We measured the lengths of newly-formed GFP^+^ sheaths in 30 µm sections of the same brains. The length-frequency distribution was shifted to shorter lengths in *Mbp^(fl/fl)^* brains compared to *Mpb^(fl/+)^*controls (**G**) and the average sheath length in *Mbp^(fl/fl)^* was around half that of controls (∼20 µm vs ∼37 µm) (**H**). (**I**) We visualized nodal structures associated with GFP^+^ myelin sheaths in individual newly-formed OLs in the motor cortex by immunolabelling with anti-CASPR together with a pan-NaV antibody. Many new sheaths in the control brains terminated at one end or both in a typical node of Ranvier with a cluster of axonal NaV (blue) adjacent to a CASPR^+^ paranode(s) (red) (arrows in **I**, high-mag images). In *Mbp^(fl/fl)^* brains the new sheaths sometimes expressed CASPR^+^ paranodes although these were frequently shorter than normal (arrows in **I**), but only rarely were new sheaths associated with clustered NaV. (**J**) The fraction of newly-formed GFP^+^ sheaths with CASPR^+^ paranodes was decreased by ∼46% in *Mbp^(fl/fl)^* brains relative to *Mpb^(fl/+)^*controls, although this did not reach statistical significance (p > 0.05). (**K**) The fraction of new sheaths that was associated with clustered NaV was reduced by ∼78% in *Mbp^(fl/fl)^* brains. *Scale bars:* (**B**) 500 µm (left), 200 µm (right and center), (**E**) 10 µm, (**I**) 20 µm.

As an additional test of *Mbp^fl^* recombination, *Ng2-Mbp-cKO* mice were combined with the *Tau-mGFP* reporter (encoding a membrane-associated GFP variant)^23^ to reveal the full morphology of OLs including their myelin sheaths.^24^ We administered tamoxifen to young adult *Ng2-Mbp-cKO: Tau-mGFP* mice (from P60–P63) and visualized myelin sheaths by co-immunolabelling for GFP and MBP at P94 (**Fig. 2D, E**). Although OLs lack the ability to form compact myelin sheaths in the absence of MBP, OLs are able to contact axons and form a few loose concentric wraps. We found a ∼74% reduction in the number of MBP^+^, GFP^+^ newly-generated sheaths compared with littermate controls (32% ± 4% of GFP^+^ sheaths in *Mbp^(fl/+)^* vs 8% ± 3% of GFP^+^ sheaths in *Mbp^(fl/fl)^*, n = 6 or 4 mice, respectively) (**Fig. 2F**). Taken together, our two independent measures indicate a rate of bi-allelic recombination in the region of 70%–75%. Myelinating OLs that were formed during development, before tamoxifen administration, are MBP^+^ but GFP-negative because differentiated OLs do not express *Ng2-CreER^TM^*.

We measured the lengths of newly-formed (GFP*-*labelled) myelin sheaths in *Ng2-Mbp-cKO: Tau-mGFP* mice and control littermates on P91. The sheath length distribution was shifted to shorter lengths in *Mbp^(fl/fl)^* compared with *Mbp^(fl/+)^* littermates (mean sheath length 36.9 ± 0.9 µm in *Mbp^(fl/fl)^*vs 19.6 ± 0.9 µm in *Mbp^(fl/+)^*, *p* <0.0001; n = 4 mice of both genotypes) (**Fig. 2G, H**). This is consistent with the observation that *Mbp* enhancer mutations that reduce MBP expression in OLs result in shorter myelin sheaths.^25^

We asked whether newly-formed MBP-negative OLs in *Ng2-Mbp-cKO: Tau-mGFP* mice can form functional nodes of Ranvier. One month post-tam, many newly-formed GFP^+^, MBP-negative myelin sheaths either lacked paranodes or had abnormally short paranodes, judged by CASPR1 immunolabelling (**Fig. 2I, J**). This likely reflects a reduced number of membrane wraps and, consequently, reduced adhesion to axons. There was also a substantial decrease in the numbers of new GFP^+^ myelin sheaths that clustered voltage-gated sodium channels, detected with a pan-NaV antibody (fraction of new sheaths with clustered NaV: 37.1% ± 1.7% in *Mbp^(fl/fl)^* vs 8.2% ± 3.9% in *Mbp^(fl/+)^*, *p* = 0.009; n = 4 and 3 mice, respectively; **Fig. 2I, K**). This suggests that without MBP, newly formed sheaths cannot establish normal nodes of Ranvier with the ability to accelerate action potentials, which is consistent with the impaired node formation observed in *shiverer* mice.^19,20^

We asked whether *Ng2-Mbp-cKO* mice could learn to run at speed on the complex wheel. We administered tamoxifen on P60–P63 and allowed a 21-day recovery period before introducing them to the complex wheel from P84–P91 (**Fig. 3A**). The *Ng2-Mbp-cKO* mice were able to master the complex wheel on a trajectory indistinguishable from normal littermate controls (**Fig. 3B**), suggesting that de novo formation of compact myelin is not critical for motor learning. We repeated this same experimental protocol using *Pdgfra-CreER^T2^* instead of *Ng2-CreER^TM^* to delete *Mbp^fl^*. These Cre drivers should be interchangeable, since both *Ng2* and *Pdgfra* are expressed predominantly in OPCs in the CNS. Indeed, we obtained the same result with both Cre drivers – *Pdgfra-Mbp-cKOs,* like *Ng2-Mbp-cKOs*, mastered the complex wheel just as well as littermate controls (**supplementary Fig. S1**).

**Figure 3.**
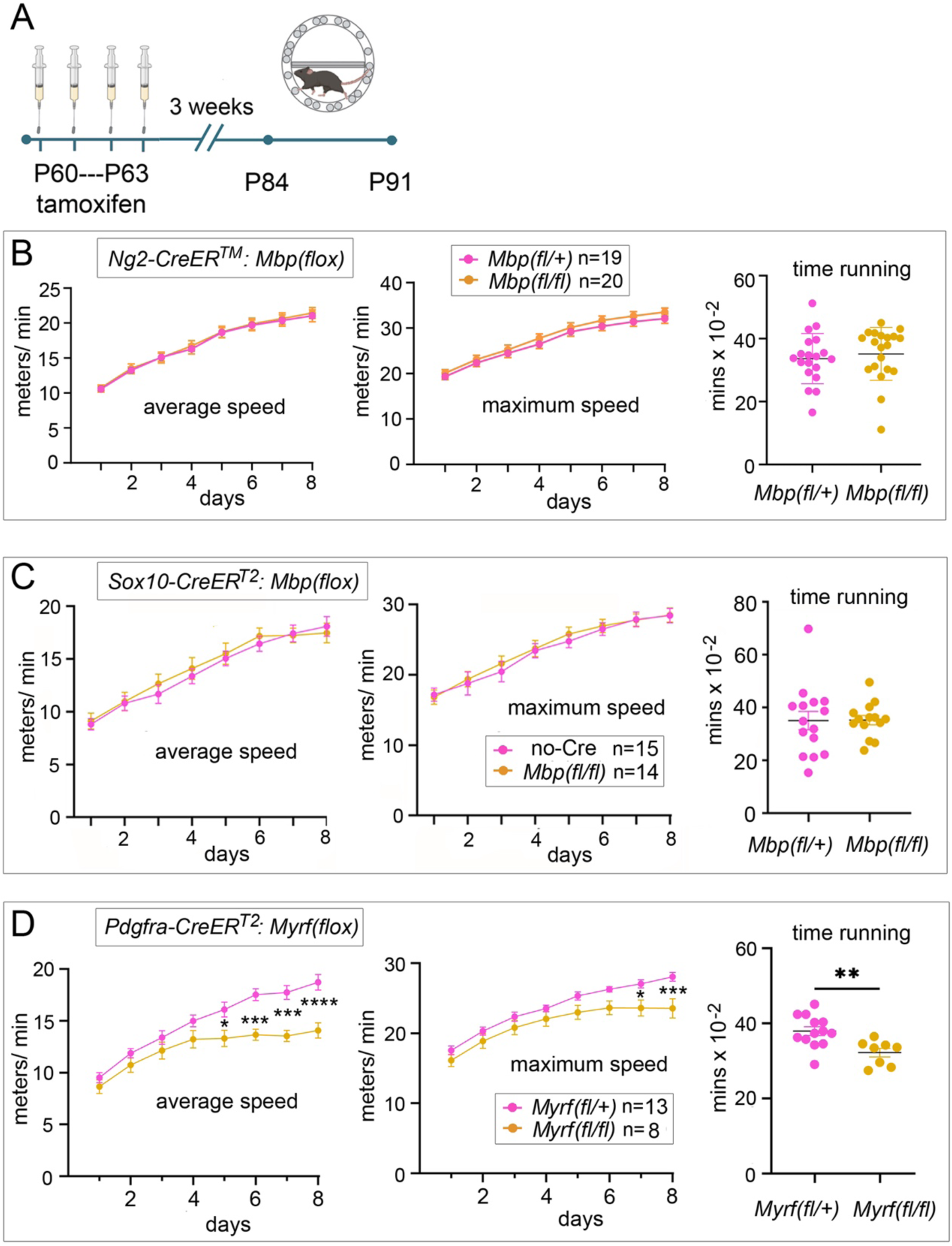
Learning to run on the complex wheel requires generation of new OLs but not compact myelin. (**A**) Timeline of the complex wheel-running experiments. Average and maximum running speeds were recorded continuously during the active period on P84–P91 inclusive. (**B**) There was no detectable difference in average or maximum speeds of *Ng2-Mbp-cKO* mice compared to control *Mbp^(fl/+)^* littermates over the 8 days running period, and no difference in time spent turning the wheel. Similar results were obtained with *Pdgfra-Mbp-cKO* mice (**supplementary Fig. S1**). (**C**) There were no differences in average or maximum speeds of *S10-Mbp-cKO* mice compared to “no-Cre” controls, and no difference in time spent turning the wheel. (**D**) In contrast, *Pdgfra-Myrf-cKO* mice were impaired in their ability to master the complex wheel compared to *Myrf ^(fl/+)^* littermate controls, and they also spent less time engaged with the wheel. * p≤ 0.05, ** p≤ 0.01, *** p≤ 0.001, **** p≤ 0.0001 (2-way ANOVA with Bonferroni’s post-hoc correction).

The incomplete recombination of *Mbp^(fl/fl)^* in *Ng2-Mbp-cKO* mice left open the possibility that the ∼25-30% of newly-formed OLs that escaped bi-allelic *Mbp* recombination, hence were still capable of MBP synthesis and myelin compaction, might be sufficient to support learning. Therefore, we switched to using *Sox10-iCreER^T2^*which, in our experience, drives close to 100% recombination of *floxed* genes in OL lineage cells, because of a very high transcription rate from the *Sox10* promoter (see Methods). *Sox10-iCreER^T2^* drives recombination at all stages of OL lineage development, unlike the OLP-specific recombination expected of *Ng2-CreER^TM^.* Hence, in adult *S10-Mbp-cKO* mice, *Mbp* transcription should be eliminated post-tam, not only in newly-differentiating OLs but also in pre-existing mature, myelinating OLs. We visualized *Mbp* mRNA and MBP protein in sections of *S10-Mbp-cKO* forebrain by in situ hybridization (ISH) and immunohistochemistry, respectively, at different times post-tamoxifen, from 2 weeks to 3 months (**Fig. 4A**). The intensity of the *Mbp* ISH signal in the corpus callosum was reduced to near-background levels relative to control littermates within 2 weeks post-tam and did not recover subsequently (**Fig. 4B**). It follows that little or no new MBP protein or compact myelin can be synthesized after this time.

**Figure 4.**
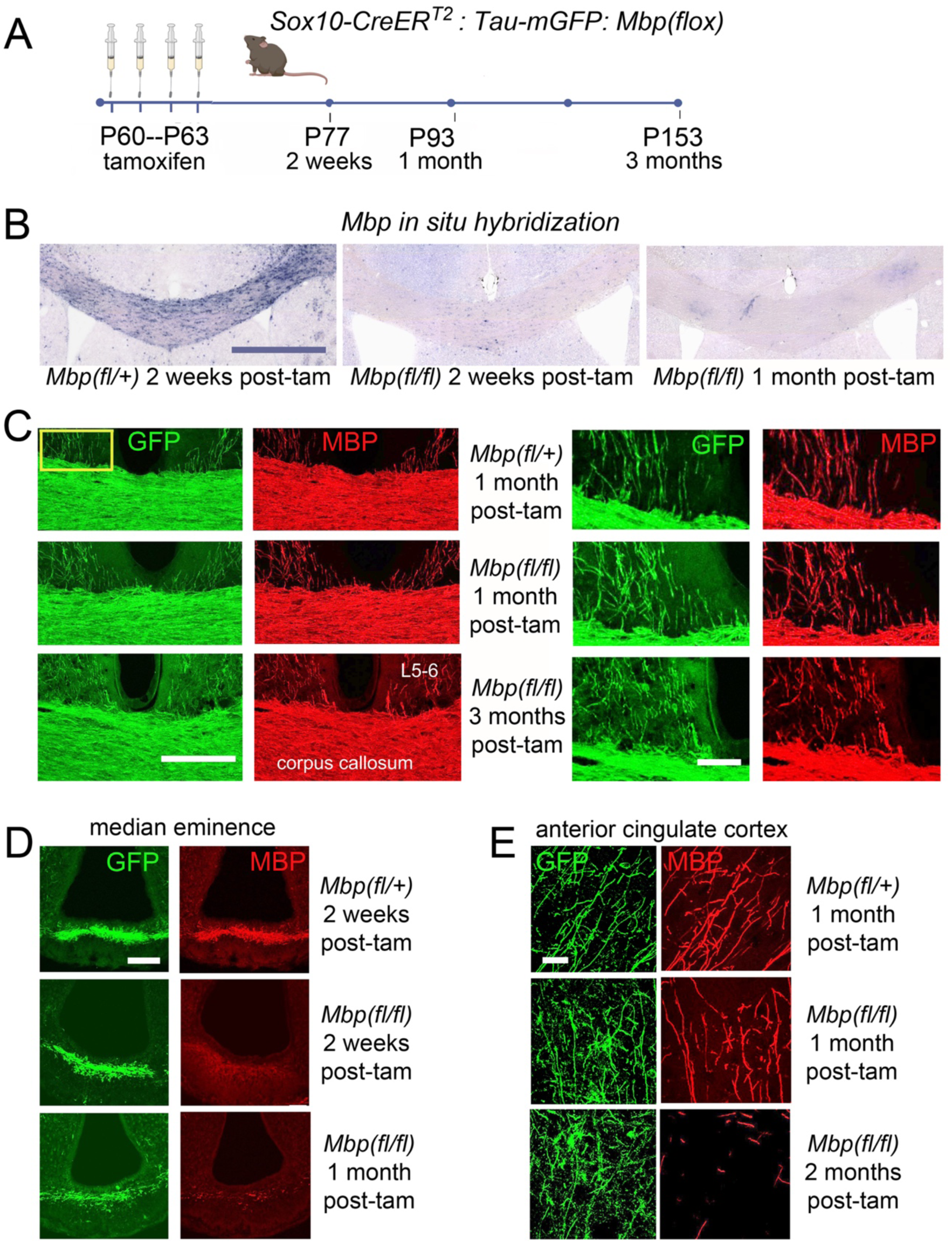
Efficient *Mbp* transcriptional block but longer-term stability of MBP protein in *S10-Mbp-cKO* mice. (**A**) Experimental protocol. *Sox10-CreER^T2^ ^(tg/+)^: Tau-mGFP^(tg/+)^: Mbp^(fl/fl)^* mice were interbred to produce *S10-Mbp-cKO* offspring together with *Sox10-CreER^T2^ ^(+/+)^*littermates (*Mbp^(fl/+)^* and *Mbp^(fl/+)^* controls). Tamoxifen was administered from P60–P63 and mice were analyzed 2 weeks, 1 month or 3 months later. (**B**) *Mbp* transcripts were detected by FISH in coronal forebrain sections of *S10-Mbp-cKO* and control brains at the level of the motor cortex at 2 weeks or 1 month post-tam. *Mbp* signal was reduced dramatically relative to controls at 2 weeks post-tam, and to background levels by 1 month. (**C**) Double-immunolabelling for GFP (green) and MBP protein (red) in coronal forebrain sections of *S10-Mbp-cKO* and *Mbp^(fl/+)^* control brains at 1 month and 3 months post-tam. The intensity of MBP fluorescence in the corpus callosum was not obviously decreased even at 3 months post-tam (left images). Using ImageJ, the MBP/ GFP intensity ratio was estimated to be reduced by ∼11% at 1 month and ∼20% at 3 months post-tam (data not shown). Higher-magnification images of the area indicated (*yellow box*) are shown on the right. Individual descending axons of cortical projection neurons are labelled for GFP and MBP; the patterns of GFP and MBP labelling are near-identical at 1 month post-tam and still very similar at 3 months, indicating that MBP is stable for at least that long in many projecting axons. (**D**) In contrast, MBP immunolabelling in the median eminence (ME) decays substantially within 2 weeks post-tam and is almost gone by 1 month. (**E**) In the anterior cingulate cortex most MBP protein survives between 1 month and 2 months post-tam. *Scale bars:* (**B**) 1 mm, (**C**) 500 µm (left) and 100 µm (right), (**D**, **E**) 20 µm.

We immunolabelled *S10-Mbp-cKO: Tau-mGFP* mice at different times post-tam for GFP (to visualize all differentiated OLs) and for MBP (to monitor the decay of MBP protein) (**Fig. 4C**). MBP immunofluorescence labelling intensity in the subcortical white matter underlying the motor cortex did not decline visibly relative to GFP over the 3 months post-tam (**Fig. 4C**). We measured MBP fluorescence intensities in *Mbp-cKOs* relative to control littermates using ImageJ and estimated that the fluorescence signal dropped ∼11% in the first 2 months post-tam and ∼20% by 3 months (data not shown).

Myelin that ensheathed descending axons at the lateral edges of the motor cortex (layers 5/6) was labelled permanently by the *Tau-mGFP* reporter post-tam, so comparison between GFP and MBP immunofluorescence labelling at different times post-tam gave an impression of the decay rate of MBP protein in cortical myelin. Even at 3 months post-tam, the majority of GFP^+^ sheaths were still clearly MBP^+^, indicating that MBP half-life was greater than 3 months in motor cortex, similar to subcortical white matter (**Fig. 4C**). This is consistent with a report that MBP is one of the most stable of all proteins in rats.^26^ This was not true of all brain regions. In the median eminence (ME, ventral hypothalamus), MBP fluorescence intensity declined noticeably relative to GFP within 2 weeks post-tam (**Fig. 4D**), supporting a previous report of rapid myelin (and OL) turnover in the ME.^27^ In the anterior cingulate cortex, MBP fluorescence decayed markedly between 1 month and 2 months post-tam (**Fig. 4E**). Meschkat et al.^17^ estimated the average half-life of MBP protein across the entire adult mouse brain to be ∼11 weeks, consistent with our observations. These data suggest that pre-existing myelin sheaths (those already in place pre-tam) persist and might function normally for many weeks post-tam, at least in the motor cortex and sub-cortical white matter.

We tested the ability of *S10-Mbp-cKO* mice to master the complex running wheel at 3 weeks post-tam, when no new MBP protein or compact myelin can be synthesized, judging by the near-complete absence of *Mbp* mRNA, yet *pre-formed* myelin can continue to function normally because of the long-term perdurance of MBP protein. Their running performance over the 8 days’ exposure to the wheel was indistinguishable from that of littermate controls (**Fig. 3C**). This result, together with our data from *Ng2-Mbp-cKOs* and *Pdgfra-Mbp-cKOs* described above, leads us to the conclusion that compact myelin formation by newly-differentiated OLs is not important for motor skill learning on the complex wheel.

*Blocking OL production in Myrf-cKO mice inhibits motor learning on the complex wheel* As a positive control for the experiments described above, we repeated one of the core experiments of McKenzie et al.,^1^ who demonstrated that *Pdgfra-Myrf-cKO* mice have a learning deficit on the complex running wheel, pointing to a requirement for newly-formed myelinating OLs in motor skill learning. That original finding has subsequently been confirmed in independent trials, using a separate *Pdgfra-Myrf-cKO* mouse colony at a different location.^9^ Nevertheless, we repeated the experiments again in our own mouse facility and reaffirmed that *Pdgfra-Myrf-cKOs* fail to learn to run at speed on the complex wheel relative to control littermates (**Fig. 3D**). We also observed that *Pdgfra-Myrf-cKOs* spent less time running on the complex wheel (**Fig. 3D**), as reported by Kaller et al., ^9^ possibly hinting at reduced motivation.

*Loss of Mct1 expression from myelinating OLs during adulthood does not inhibit skill learning* In addition to increasing the speed of propagation of APs, myelinating OLs are believed to support neuronal metabolism by transferring substrates for ATP production (e.g. lactate) into their partner axons via the peri-axonal space, through monocarboxylate transporters MCT1 and MCT2 in the apposed myelin and axonal membranes, respectively ^28–30^. This would be expected to boost energy production in axons, e.g. to fuel regeneration of the resting membrane potential at nodes of Ranvier during rapid firing. We asked whether metabolic coupling between newly-formed OLs and their newly-ensheathed axons is important in motor learning. We generated *Pdgfra-CreER^T2^*: *Mct1^(fl/fl)^* (*Pdgfra-Mct1-cKO*) and *Sox10-CreER^T2^: Mct1^(fl/fl)^* (*S10-Mct1-cKO*) mice, administered tamoxifen in young adulthood (P60–P63) and tested their ability to master the complex running wheel 3 weeks post-tam. We found no difference in running ability of either *Pdgfra-Mct1-cKO* or *S10-Mct1-cKO* mice on the complex wheel relative to control littermates (**supplementary Fig. S2**). Thus, MCT1-mediated lactate transport from newly-differentiated OLs appears not to be critical for motor skill learning. However, we were unable to detect MCT1 immunoreactivity above background in bona fide OLs or their myelin sheaths in the grey or white matter of control mice, despite being able to detect MCT1 readily in brain capillary endothelial cells. Therefore, we were unable to demonstrate that *Mct1* was efficiently deleted in these experiments – although we have no reason to think it was not (see Discussion).

### Neo-myelination is not required for long-term consolidation of motor or fear memories

We asked whether MBP synthesis and myelination might be necessary to consolidate long-term motor memory. We allowed *Ng2-Mbp-cKO* mice and control littermates to self-train on the complex wheel for 8 days starting 3 weeks post-tamoxifen, removed the mice to their home cages for 30 days, then re-introduced them to the complex wheel (**Fig. 5A**). After reintroduction, there was no discernible difference between the performances of *Ng2-Mbp-cKO* mice and littermate controls; within 4 days both groups regained and exceeded the average and maximum running speeds they had attained during their initial 8 days of training (**Fig. 5B**). It appears, therefore, that formation of compact myelin and nodes of Ranvier by newly-differentiating OLs is not required either for initial performance improvement on the complex wheel (motor learning) or for 30 day retention of the learned skill (motor memory consolidation).

**Figure 5.**
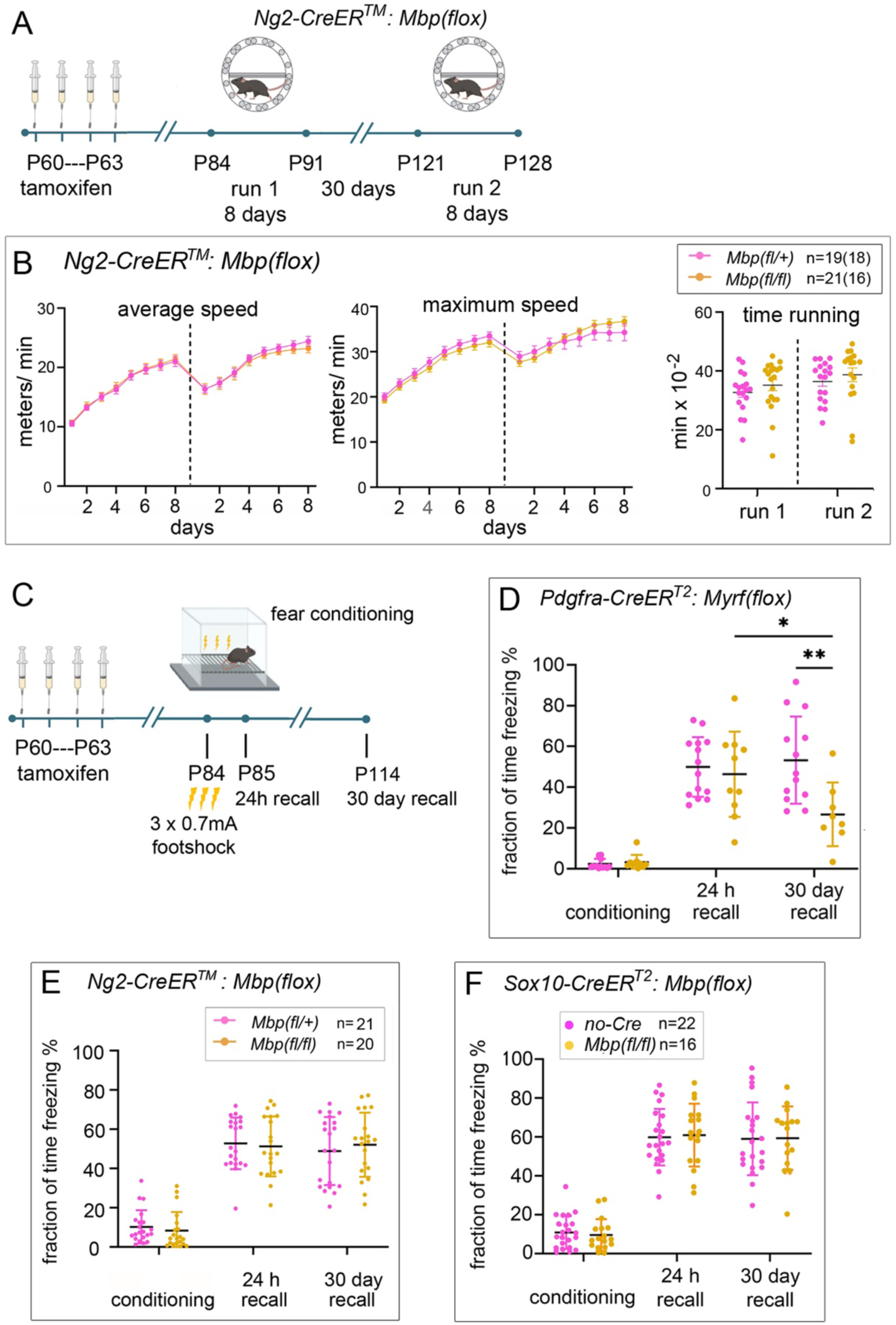
Long-term consolidation of motor or fear memories does not depend on de novo compact myelin formation. **(A)** We asked whether *Ng2-Mbp-cKO* mice that had self-trained on the complex wheel for 8 days (“run 1”) retained their learned skill after being removed to their home cage for 30 days then re-introduced to the wheel for a second 8-day period (“run 2”). (**B**) During both run 1 and run 2, the running performances of *Ng2-Mbp-cKO* mice and control *Mbp^(fl/+)^*littermates were indistinguishable. Their average and maximum speeds on the first day of run 2 were around half what they were on the final day of run 1, then their performance improved during run 2 to slightly exceed that achieved on the final day of run 1. Both groups spent similar times engaged with the wheel (“time running”). Note that some mice from both cohorts were not used for run 2, so “n” was lower for run 2 than for run 1 (18 vs 19 for *Mbp^(fl/+)^*controls and 16 vs 21 for *Ng2-Mbp-cKO*). (**C**) We also tested the fear responses of mice during contextual fear-conditioning, and their subsequent short-term and long-term (remote) fear memories. (**D**) As a control, we confirmed previous reports that *Pdgfra-Myrf-cKO* mice display normal fear-conditioning responses and normal short-term (24 h) recall, but defective long-term (30 day) recall. * p≤ 0.05, ** p≤ 0.01 (2-way ANOVA with Bonferroni’s post-hoc correction). (**E**, **F**) In contrast, both *Ng2-Mbp*-cKO (**E**) and *S10-Mbp-cKO* (**F**) were indistinguishable from control littermates at all time points including 30 days post-conditioning. These data indicate that blocking de novo compact myelin formation is dispensable for consolidation of fear or motor memories.

Previously it was reported that blocking new OL generation in *Ng2-Myrf-cKOs*^7,8^ inhibits remote (30 day) recall of contextual fear memories. Blocking OL generation in *Ng2-Myrf-cKOs* or *Pdgfra-Myrf-cKOs* also inhibited remote recall of spatial memory in the Morris water maze.^8^ These results suggested that new myelin formation is important during consolidation of fear and spatial memories. We confirmed the requirement for newly-generated OLs in consolidation of contextual fear memory. Three weeks after tamoxifen administration, *Pdgfra-Myrf-cKOs* and control littermates were placed in a fear-conditioning chamber for 5 min and subjected to three mild foot shocks, before returning them to their home cages. After 24 h, the mice were reintroduced to the chamber for 5 min without foot shocks and freezing behaviour was recorded (**Fig. 5C**). The same test was repeated 30 days after conditioning to assess long-term memory retention. Control mice spent most of the session freezing at both 24 h and 30 day points, indicating successful fear memory formation and long-term consolidation (**Fig. 5D**). In contrast, *Pdgfra-Myrf-cKO*s showed normal freezing behaviour 24 h after conditioning but significantly reduced freezing after 30 days (**Fig. 5D**), confirming that new OL formation is required for fear memory consolidation, as previously reported.^7,8^ However, in the absence of new MBP synthesis from newly-formed OLs (*Ng2-Mbp-cKO*) or all OL-lineage cells (*S10-Mbp-cKO*), mice displayed normal freezing behaviour at both 24 h and 30 day time points (**Fig. 5E**, **F**). Thus, compact myelin and node formation by newly-differentiating OLs are not essential for long-term fear memory consolidation.

*MCT1 expression by newly-formed OLs is dispensable for consolidation of motor or fear memories* We tested the ability of *Pdgfra-Mct1-cKOs* and *S10-Mct1-cKOs* to form and consolidate long-term fear memories. In the absence of MCT1 in newly-formed OLs (*Pdgfra-Mct1-cKO*) or in all OL lineage cells (*S10-Mct1-cKO*), mice displayed normal freezing behaviour at both 24 h and 30 day time points (**supplementary Fig. S2**). Therefore, MCT1-mediated lactate transport by newly-differentiated OLs also appears not to be essential for long-term fear memory consolidation.

## Discussion

It has been established previously (and confirmed here) that conditional knockout of *Myrf,* aimed at blocking de novo production of myelin-forming OLs during adulthood, limits the ability of mice to improve their performance through training (i.e. to learn), both in a gross motor task (complex running wheel)^1,2,9^ and in cognitive tasks in a T-maze or 8-arm radial maze.^10^ *Myrf-cKO* also inhibits long-term consolidation of spatial or contextual fear memories, without affecting spatial or fear learning per se.^7–9^ These and other studies have suggested that that de novo myelination is needed for some forms of learning as well as for long-term memory consolidation. Against this trend, Wang et al.^6^ found that *Myrf-cKO* mice became *more* adept than littermate controls – not less – at a task involving reaching/ grasping for food pellets. However, consolidation of the learned skill was impaired when *Myrf-cKO* was induced after learning. ^6^

We have now shown that conditional block of MBP synthesis during adulthood, which prevents newly-differentiating OLs from forming compact myelin sheaths, does not inhibit either motor skill learning on the complex wheel or contextual fear learning, nor does it prevent consolidation of those learning experiences into long-term motor or fear memories. This was not our expectation, given the *Myrf-cKO* studies outlined above, and leads us to alter the way we think about the role of OL lineage cells in learning and memory processes.

### How do OLs contribute to mechanisms of learning and memory?

#### Canonical role of myelin in speeding action potentials

Based on a variety of experimental approaches, we and others have speculated that an important role for newly-generated OLs might be to detect and myelinate axons that fire during learning behaviours, increasing the speed and amplitude of action potentials (APs) in more active circuits over those in less-active or silent parallel circuits (“adaptive myelination”).^1,4,5^ This would frame learning as a kind of natural selection at the level of neural circuits.^1^ A related suggestion has been that adaptive myelination fine-tunes transmission times between widely separated brain regions or within a given brain region, leading to closer coherence of rhythmic firing patterns between or within those regions, a phenomenon that is known to accompany learning and memory formation.^38–40^ For example, convergence of so-called theta rhythms (4-12 Hz oscillations) in hippocampal CA1 and medial prefrontal cortex has been shown to correlate with a correct choice of goal-arm, but not an incorrect choice, in an “H-maze” test of spatial working memory in rats.^41^ In keeping with this idea, *Myrf-cKO* mice were shown to have impaired temporal coordination between high-frequency bursts (“ripples”) in the hippocampus and low-frequency oscillations (“spindles”) in thalamo-cortical circuits (known as ripple-spindle coupling).^8,42^ *Myrf-cKO* mice also had increased electro-encephalogram (EEG) spectral power, again linking OL genesis to oscillatory brain activity. ^9^

These ideas, based on the canonical role of myelinating OLs in controlling AP speed, seem at odds with our present findings from *Mbp-cKO* mice. However, uncompacted “*shiverer-*type” membrane wraps, which also form in our *Mbp-cKO* mice, can still exert a positive effect on conduction. Abnormal nodes or hemi-nodes, some containing clustered NaV, can still form in association with *shiverer-*type loose myelin wraps (references 19, 20 and this paper) and are able to support AP propagation at speeds intermediate between those observed in fully myelinated versus unmyelinated axons in wild type mice (one-third to one-half the speed of normally myelinated axons in spinal cord).^20,43^ Other functional differences between wild type and *shiverer-*type myelinated axons are that the amplitudes of APs are significantly reduced in the latter, and their ability to sustain high-frequency repetitive firing is greatly reduced.^20,43^ Therefore, while it remains a formal possibility that loose *shiverer-*type myelin generated de novo in *Mbp-cKO* mice might contribute to learning by increasing AP conduction speed, our failure to detect even a slight reduction in the learning ability of *Mbp-cKOs* prompts us to consider alternative possibilities.

#### Metabolic support of axons

An alternative role for newly-formed OLs in learning might be to support the metabolism of newly-myelinated axons. For example, it has been reported that myelinating OLs release glycolytic products lactate and/or pyruvate into the periaxonal space through MCT1, from where they can enter the ensheathed axons through MCT2 to feed oxidative phosphorylation for ATP production.^28—30^ We asked whether this form of metabolic coupling is important during learning; we generated *Mct1-cKOs*, using *Pdgfra-CreER^T2^* or *Sox10-CreER^T2^* as drivers, but we observed no motor learning deficits on the complex wheel (**supplementary Fig. S2**). This might be 1) because metabolic coupling via MCT1 is unimportant for learning, or 2) because we failed to delete *Mct1^flox^* effectively. We could not quantify recombination efficiency because we could not detect MCT1 expression in OLs by immunolabelling, even before tamoxifen administration. However, we believe that *Mct1* is likely to be deleted efficiently in OL lineage cells because of our experience that *Sox10-CreER^T2^* drives very efficient recombination of other *floxed* alleles including *Myrf ^fl^* and *Mbp^fl^*, and there is no reason to think that *Mct1^fl^* should be different because the distance between its *loxP* sites (<2.5 kb)^44^ is well within the limit for high-efficiency excision (<4 kb).^45^ Moreover, Philips et al.^46^ found no acute learning deficits of constitutive *Sox10-Cre: Mct1 ^(fl/fl)^* mice in passive avoidance, Y-maze or fear-conditioning tests, although late-onset axonal pathology developed after 8 months. The combined evidence therefore suggests that MCT1 in OL lineage cells is not required to support learning and memory.

Even if MCT1 is not involved, we cannot rule out that some form of metabolic coupling might be important for learning and memory, because there are other molecules or pathways through which newly-formed OLs might transfer energy substrates into the periaxonal space. These include, for example, another member(s) of the MCT family, exosome release and reuptake ^47^, or CONNEXIN-47 hemichannels.^46^ There is also recent evidence that, under severe energy demand, myelin membranes themselves might be used as an accessible source of lipids that can feed into oxidative phosphorylation and ATP synthesis through fatty acid beta-oxidation.^48^

### Pre-myelinating OLs in learning and memory?

We found that adult-born CC1^+^ OLs are formed, survive and accumulate normally in *Mbp-KO* mice, whereas OLP differentiation is interrupted at an earlier stage in *Myrf-KOs* such that newly-formed CC1^+^ OLs are almost completely absent.^1^ This raises the possibility that an early pre-myelinating stage of the OL lineage (pmOLs), rather than mature myelinating OLs, are critical for learning.

Mice start to improve their performance on the complex wheel very quickly, within the first 2 h of introduction to the wheel.^2^ Correspondingly, the running performance of *Pdgfra-Myrf-cKO* mice falls behind that of their control littermates within the same short time frame of ∼2 h.^2^ This does not seem to allow time for OL differentiation and the elaboration of mature myelin sheaths, which in zebrafish takes one or two days, although axon selection and the initiation of myelination can occur within a few hours.^49^ Moreover, in a radial maze working memory task, success rate correlates more closely with the proportion of OLPs that is induced to enter the cell cycle (PDGFRA^+^, EdU^+^/ PDGFRA^+^) than with the number of new OLs (CC1^+^, EdU^+^) that form during the learning period (R^2^ = 0.84 vs 0.61, respectively, in corpus callosum; R^2^ = 0.72 vs 0.21 in anterior cingulate cortex).^10^ These data also hint that early stages of the OL lineage (OLPs or pmOLs) are more important in determining learning outcomes than mature, fully-myelinating OLs.

OLPs have been implicated in axonal pruning in the developing mouse cerebral cortex^50^ and the developing zebrafish optic tectum.^51^ There is also evidence that OLPs can phagocytose neuron-neuron synapses in the mouse visual cortex during the critical period of synaptic refinement (P20-P27) and that their phagocytic capacity can be modified by experience (exposure of dark-reared mice to light).^52^ This raises the possibility that OLPs might perform a similar function during experience-dependent plasticity in the adult brain. However, it is not obvious how *Myrf-cKO* might impact this pruning function of OLPs, since OLPs do not express *Myrf* and OLP numbers are not altered substantially by *Myrf-cKO.*^1^ Since Auguste et al.^52^ found that phagocytic activity was not uniform across their OLP population – about 5% of the population was >10 times more active than the majority – it is conceivable that at least some of the synaptic pruning ascribed to OLPs might actually be a specialization of early-differentiating OLs.

### OL dynamics and the central control of energy balance

There is accumulating evidence that OL lineage dynamics in the median eminence (ME) in the basal hypothalamus are regulated by nutritional signals and play a key role in the central control of energy balance, together with hypothalamic neurons of the arcuate nucleus (ARC). There is an unusually high turnover rate of OL lineage cells in the ME of adult mice, and OLP proliferation and OL differentiation/ survival are positively regulated by food intake.^27,53^ Receptors for the adipose-derived hormone leptin, which regulates body fat mass by controlling appetite and satiety, are concentrated on the dendrites of ARC neurons. OLPs are in contact with these dendrites and appear to be required for their maintenance.^54^ Hence, drug- or radiation-mediated ablation of OLPs, either brain-wide or targeted to the ME, resulted in leptin insensitivity and weight gain in mice.^54^ Preventing new OL generation in *Pdgfra-Myrf-cKO* mice resulted in reduced food intake accompanied by reduced energy expenditure, including reduced core body temperature and reduced ambulatory activity.^27^

These systemic effects of *Myrf-cKO* could potentially contribute indirectly to the learning and memory deficits that we and others have described. However, since learning modulates OLP proliferation and OL differentiation in brain regions relevant to the learned behaviour in wild type mice, and because the scale of OL dynamics in those regions correlates with performance in the learned task but not with food intake,^5,6,10^ we think that perturbation of local OL dynamics is more likely to be responsible for the learning/memory phenotypes of *Myrf-cKOs* than systemic effects originating in the ME. Nevertheless, we should keep in mind that systemic effects of *Myrf-cKO* could potentially influence the outputs of “appetitive” learning paradigms, in which dietary restriction is employed to motivate reward-seeking behaviour – especially in view of a recent report that OPC proliferation is glucose-dependent.^55^

### Role of extracellular matrix in learning and memory

There is increasing interest in the properties and functions of distinct classes of extracellular matrix (ECM)-rich structures in the brain, including perineuronal nets (PNNs) ^56^, perinodal ECM,^56^ and dandelion clock-like structures (DACS).^57,58^ There is evidence of functional connections among these ECM structures, OL lineage cells and neural circuit plasticity,^59,60^ so it is plausible that a key function of OLPs and/or pmOLs in learning and memory might be to lay down or modify ECM components in in the brain. This would be consistent with the contrasting results of our behavioural experiments with *Mbp-cKO* and *Myrf-cKO* mice reported here.

In the ME, for example, PNNs envelop ARC neurons and have been postulated to regulate their exposure to blood-borne nutritional signals, thereby controlling feeding behaviour. ^53,61^ OLPs and newly-differentiating pmOLs express several chondroitin sulphate proteoglycans (CSPGs) that are components of PNNs; pmOLs also express enzymes such as ADAMTS4, a metalloproteinase that can cleave CSPG core proteins. Hence, pmOLs might regulate ARC neuron activity by laying down and modifying PNNs in the ME. Consistent with this idea, ADAMTS4 is down-regulated in OLs in the ME during fasting, and up-regulated again following refeeding.^53^

PNNs are also prominent in other brain regions, particularly cortical regions including motor and prefrontal areas, where they are mainly associated with parvalbumin-expressing inhibitory interneurons. There is a growing literature on the role of PNNs in neural circuit plasticity, e.g. in synaptogenesis and closure of critical periods, memory and cognitive processes and various psychiatric and psychosocial conditions.^62,63^ Xin et al.^64^ showed that blocking OL neogenesis in adolescent (P14) *Ng2-Myrf-cKO* mice reduced inhibitory transmission in the visual cortex subsequently, although the distribution and labelling intensity of PNNs was not perturbed.

DACS were originally thought to be a product of astrocytes^57^ but have now been shown to correspond to the outlines of newly-differentiating pmOLs and the “ghosts” of pmOLs that that undergo apoptotic dell death^58^ as a result of homeostatic cell number control.^24,58,65,66^ DACS persist for up to 10 days even after death of the pmOLs that produced them, so they accumulate in brain tissue and could comprise a significant fraction of the ECM in the adult mouse cortex.^60^ Therefore, it will be interesting to discover in future whether pmOL-derived DACS play a role in learning and memory.

## Conclusion

We have provided evidence that the role of newly-differentiating OLs in neural circuit plasticity does not rely on the most obvious and best-understood property of OLs – that is, to form compact myelin around axons to accelerate their conduction speeds. Instead, it is more likely to involve some ancillary role of newly-generated, pre-myelinating OLs. What this key non-canonical function of OLs might be is currently unknown, but some possibilities have been discussed above.

Distinguishing among these and other functions of OLs will require further genetic manipulation of the OL lineage coupled to behavioural and physiological readouts. This is an important area for further investigation because of its wide-ranging implications, not only for cognition and memory, but also for normal ageing processes and neuropsychiatric and psychosocial disorders.

## Methods

### Mice

Mouse experiments were pre-approved by the UCL Animal Welfare and Ethical Research Board and conformed to the Animals (Scientific Procedures) Act 1986 of the UK Government and its subsequent amendments. Mice were maintained on an artificial 12 h light-dark cycle, commencing at 7 am. Male and female mice were used in roughly equal proportions in all experiments.

We designed the *Mbp^fl^* target vector and provided the design to Cyagen (Santa Clara, California) to custom-make *Mbp^fl^* mice. Exon 5 of the *Golli-Mbp* gene on mouse chromosome 18 was flanked by *loxP* sites inserted into intron 4 and exon 5c such that Cre recombination deletes 1531 bp (the cKO region) including the classical *Mbp* transcriptional start site in exon 5b (Fig. 1A). To engineer the targeting vector, the cKO region and flanking regions (homologous arms) were PCR-amplified from clones RP24-285N19 and RP24-149A18 (Roswell Park Cancer Institute C57BL/6J male mouse BAC library). The *Neo^R^*cassette was flanked by *frt* self-deleting anchor sequences and Diphtheria toxin (DTA) was used for negative selection in C57BL/6 embryonic stem cells.

To generate OL lineage-specific conditional knockout (cKO) offspring, *Mbp ^fl^* mice were crossed with *Ng2-CreER^TM^,*^22^ *Sox10-iCreER^T2^,*^1^ or *Pdgfra-CreER^T2^* mice ^67^ on a C57BL/6J background, to generate *Ng2-Mbp-cKOs, S10-Mbp-cKOs* or *Pdgfra-Mbp-cKOs* together with *Mbp^(fl/+)^* or “no-Cre” controls, as noted in the figures. Male *Mbp^fl^* mice were also crossed with female *Sox10-Cre* mice ^68^ to generate constitutive *Mbp-KOs. Tau-mGFP* (encoding membrane-bound GFP knocked in at the *Tau* locus)^23,24^ was included in some experiments to visualize OL lineage cells post-tamoxifen. For example, *Ng2-CreER^TM^ ^(tg/tg)^: Tau-mGFP ^(tg/+)^: Mbp^fl/+^* males and females were interbred to produce mixed litters of *Ng2-CreER^TM^ ^(tg/tg)^: Tau-mGFP ^(tg/+)^: Mbp^(fl/fl)^ cKOs* along with *Mbp^(fl/+)^*controls. *Myrf ^fl^* mice^15^ were crossed with *Pdgfra-CreER^T2^,* on a C57BL/6J background. *Mct1^fl^* mice^44^ were crossed with *Pdgfra-CreER^T2^* or *Sox10-CreER^T2^*, also on a C57BL/6J background. Genotyping was by PCR amplification of ear clip genomic DNA.

To initiate CreER-*lox* recombination, tamoxifen (Sigma-Aldrich) was administered by oral gavage on four consecutive days P60–P63, at 300 mg tamoxifen/kg body weight. Stock solution was made by mixing tamoxifen powder at 40 mg/ml with corn oil (Sigma-Aldrich) and sonicating the suspension for 1 h at 20-37°C.

### Genotyping by polymerase chain reaction

For genotyping, 25 μL PCR reactions containing 0.2 mM dNTPs (Amersham Pharmacia Biotech), 1 μl of Taq DNA polymerase and 20 pmol of each primer (**supplementary Table S1**) in PCR buffer [500 mM KCl, 1.5 mM MgCl_2_,1% (v/v) Triton X-100, 100 mM Tris-HCl pH 9.0] were incubated as follows: 94°C/ 4 min, followed by 40 cycles of (94°C/ 30 s – 60°C/ 45 s – 72°C/ 60 s), concluding with 72°C/ 10 min. PCR products were visualized using the Qiaxcel advanced capillary electrophoresis system (Qiagen).

### Histology and immunolabelling

Mice were perfused transcardially with 4% (w/v) paraformaldehyde (PFA, Sigma) in phosphate-buffered saline (PBS). Brains were removed and sliced using a cutting matrix (Agar Scientific) and stored overnight in 4% PFA; Sigma) at 4°C. This was followed by cryoprotection in diethyl pyrocarbonate (DEPC)-treated 20% (w/v) sucrose (Sigma) in PBS at 4°C until the tissue sank.

Brain tissue was embedded in Optimal Cutting Temperature (OCT) compound (Tissue-Tek), frozen on dry ice and stored at –80°C before cutting 20 µm or 30 μm coronal cryosections on a Bright OTF 5000 cryostat. Sections were stored in PBS containing 0.02% (w/v) sodium azide at 4°C for up to a week before immunolabelling. For node of Ranvier labelling with a pan-NaV antibody, mice were perfused transcardially with 9% (v/v) glyoxal (Sigma), 8% acetic acid pH 4,^69^ and processed further as above.

Sections were washed 3x 10 min in PBS, immersed in 300 µL of blocking buffer [0.1% (v/v)] Triton X-100, 10% (v/v) fetal bovine serum (FBS) in PBS] for 1 h, then incubated overnight in primary antibody (**supplementary Table S2**) in blocking buffer, 300 μL per well of a multiwell plate (all at 4°C). The following day, sections were washed 3 x 5 min in PBS before incubating for 1–2 h at 20–25°C with 300 μL Alexa Fluor-488, −568 or −647 secondary IgG (Invitrogen) in blocking buffer (1:500 or 1:1000) containing Hoechst 33258 dye (Sigma-Aldrich, 0.2 μg/ml). The sections were again washed 3 x 5 min in PBS before mounting on glass slides for microscopy.

### RNA in situ hybridization

Our ISH protocol was described previously (https://www.homepages.ucl.ac.uk/~ucbzwdr/-*> lab protocols*).^70^ Briefly, 20 µm cryosections were mounted on glass microscope slides and incubated at 65°C with fluorescein isothiocyanate (FITC)-labelled *Mbp-*riboprobes in wash buffer [50% formamide, saline sodium citrate (SSC), 0.1% (v/v) Tween20 in double-distilled H_2_O]. The following day, sections were washed 2x 30 min at 65°C in wash buffer and 3 x 10 min at 20-25°C in MABT [100 mM maleic acid, 150 mM NaCl, 0.1% (v/v) Tween-20, pH7.5]. Sections were incubated in blocking solution [MABT containing 2% blocking reagent (Roche catalogue number 1 096 176), 10% v/v heat-inactivated goat serum] for 1 h. For single-probe ISH, the FITC signal was incubated overnight at 4°C with an alkaline phosphatase (AP)-conjugated anti-FITC Fab_2_ antibody fragment (Roche Molecular Biochemicals). Sections were washed 3 x 5 min in MABT before incubating for 3 h in tyramide signal amplification (TSA) system following the manufacturer’s instructions (NENTM Life Science Products). Finally, sections were washed 3 x 5 min in MABT at 20–25°C before mounting on glass slides under coverslips.

### EdU labelling in vivo and estimation of Cre-lox recombination efficiency

To estimate *Mbp ^fl^*recombination efficiency and OLP cell cycle parameters, we treated P60 *Ng2-CreER^TM^: Mbp^(fl/fl)^:Tau-mGFP* and their *NG2-CreER^TM^: Mbp^(fl/+)^:Tau-mGFP* littermates with tamoxifen for 4 days (P60–P63), then we administered 5-ethynyl-2’-deoxyuridine (EdU) in the drinking water (0.2 µg/ml) for 10 days, starting on P80. The mice were analyzed immediately after EdU treatment by fluorescence ISH (FISH) with an *Mbp* riboprobe, followed by Click-iT detection of EdU (ThermoFisher). *Mbp^+^,* EdU^+^ cells were scored as newly-formed OLs that had escaped Cre recombination; the numbers of such OLs in *Mbp-*cKOs vs controls allowed us to estimate recombination efficiency to be ∼70% in the subcortical white matter.

Recombination efficiencies in *Sox10-Cre: Mbp^(fl/fl)^*and *Sox10-iCreER^T2^: Mbp^(fl/fl)^* mice were estimated by measuring the average intensities of *Mbp* ISH signals, relative to *Mbp^(fl/fl)^* controls, using ImageJ (sum of the gray values of all pixels divided by the number of pixels). The recombination efficiency was >90% in subcortical white matter for both *Sox10-Cre* and

### Sox10-iCreER^T2^

#### Imaging and cell counts

Immunolabelled sections were visualized in a Leica confocal microscope with LAS-AF software or a Zeiss 880 Airyscan, applying standard excitation and emission filters for DAPI (350 nm), FITC (Alexa Fluor-488), Cy3 (Alexa Fluor-568) and Far Red (Alexa Fluor-647). Z-stacks with 0.88 μm spacing were processed for further analysis using ImageJ software (Fiji).

We counted cells manually in low-magnification micrographs (x20 objective) of non-overlapping areas in coronal sections of the medial subcortical white matter, between the lateral edges of the lateral ventricles, using ImageJ. We counted three regions of interest in at least three sections from three or more mice of each genotype and age group (P14, P28, P46).

To quantify newly-generated myelin sheaths and nodes of Ranvier, we captured high-magnification fluorescence micrographs (x63 objective) of *Tau-mGFP^+^* motor cortex immunolabelled for GFP and MBP and counted (GFP^+^, MBP^+^) and (GFP^+^, MBP^-^) myelin sheaths. We scored newly-formed nodes as follows: 1) each one lay within the section at the end of a GFP^+^ myelin sheath, 2) it contacted or overlapped a CASPR1^+^ paranodal junction, 3) it ended next to a cluster of pan-NaV expression. We measured lengths of GFP^+^ myelin sheaths in 30 µm sections by selecting sheaths that terminated at both ends within the section.

#### Electron microscopy

Mice were perfusion-fixed with 2.5% (v/v) glutaraldehyde and 2% (w/v) PFA in 0.1 M sodium cacodylate buffer (pH 7.4). Brains were post-fixed by immersion in the same fixative overnight at 4°C, before transferring to 0.1 M PBS for shipment from UK at ambient temperature. In Japan, thick sagittal slices were prepared at <1 mm thickness from 4.5 x 2 mm tissue blocks containing the whole length of the subcortical white matter (SWCM). The sections were immersed in 1% (w/v) osmium tetroxide solution for 2 h at 4°C, dehydrated through a series of graded alcohols and embedded in Epon 812 resin (TAAB Laboratories, UK). Ultrathin parasagittal sections (100 nm) were cut on an ultramicrotome (Ultracut UCT, Leica) and collected on tin-coated glass slides, stained with uranyl acetate and lead citrate and imaged in a scanning EM equipped with a back-scattered electron beam detector (Hitachi SU8010) at 1.0–1.5 kV accelerating voltage. This allowed low-magnification, wide-field imaging of white and gray matter.

#### Complex running wheel

Our complex wheel running task is described by McKenzie et al.^1^ An aluminium running wheel (12.7 cm diameter) with 38 rungs was modified by removing 16 rungs in a quasi-random manner to create tandem-duplicated 11-rung patterns with unequal gaps. Mice were introduced to the complex wheel task three weeks after the last day of tamoxifen administration (allowing time for recombination and for the mice to recover from tamoxifen). Their whiskers were clipped to prevent the mice from using them to detect oncoming rungs. Each mouse spent 8 consecutive days in the wheel cage; wheel speed was monitored automatically by an infra-red detector at one-minute intervals over the active (dark) period. We recorded instantaneous and average speeds, accumulated distance run and time spent running at >1 m/min by individual mice during the active period. In some experiments, mice were re-introduced to the complex wheel for a further 8 days after a 30 day rest period in their home cage.

#### Fear conditioning

Mice were acclimatised to handling for 7 days prior to conditioning and brought into the behavioural room 1 h before conditioning. To establish a fear memory, we used a fear conditioning chamber (Actimetrics, Model 80014AT) in an isolation cubicle. Mice were introduced to the chamber for 5 min exposure to the conditioning cues (chamber, lights, white noise and scent of anise). Following that they received 3 foot-shocks of 0.7 mA x 2 s, separated by 30 s. Mice were returned to the conditioning chamber and exposed to the same cues 24 h or 30 days post-conditioning for 5 min without further electrical shock. Videos were recorded of all trials and freezing behaviour automatically calculated using FreezeFrame 6 software and plotted as a percentage of the 5 min exposure time.

#### Statistics and graphics

Prism 6.0 (GraphPad) was used for statistical analysis. Running wheel data were analyzed by calculating average speed (m/ min) for individual mice over the whole dark cycle, then mean ± s.e.m. for the whole cohort. To compare differences between cohorts at a given time point we used ANOVA with Bonferroni’s post-hoc test. Two-way ANOVA was used when time and genotype were considered as two independent variables. To test for normality of a data distribution, or whether two datasets have the same distribution, we used the 1- or 2-sample Kolmogorov-Smirnov non-parametric test. All experiments and analyses were carried out with the experimenter blind to the sex and/or genotype of the mice. Graphs were generated in GraphPad, annotated and arranged in Adobe Photoshop Elements. Other display items were created using BioRender and Photoshop Elements.

## Acknowledgements

We thank our colleagues in the Wolfson Institute for Biomedical Research for help, advice and comradeship. This work was supported by the Wellcome Trust (108726/Z/15/Z and 214286/Z/18/Z to W.D.R.), a Sanming Project (SZSM201911003) funded by the Municipal Government of Shenzhen, China to W.D.R and H.L., and a Career Development Award from the UK Medical Research Council (MR/X019977/1) to M.S.

## Author contributions

W.D.R. and H.L. conceived the project and secured funding. W.D.R., H.L., S.G.N. and M.S. designed the experiments. M.S and S.G.N. conducted most of the experiments with help from Y.J., A.H. and M.L. O.T. and K.T. provided electron microscopy analysis. T.P. and J.R. provided *Mct1^fl^* mice. W.D.R. wrote the paper with input from M.S., S.N., A.H. and H.L.

## Artificial intelligence

No artificial intelligence tools were used in the generation, analysis or reporting of the data presented here.

## Supplemental data relating to

**Supplementary Table S1.**
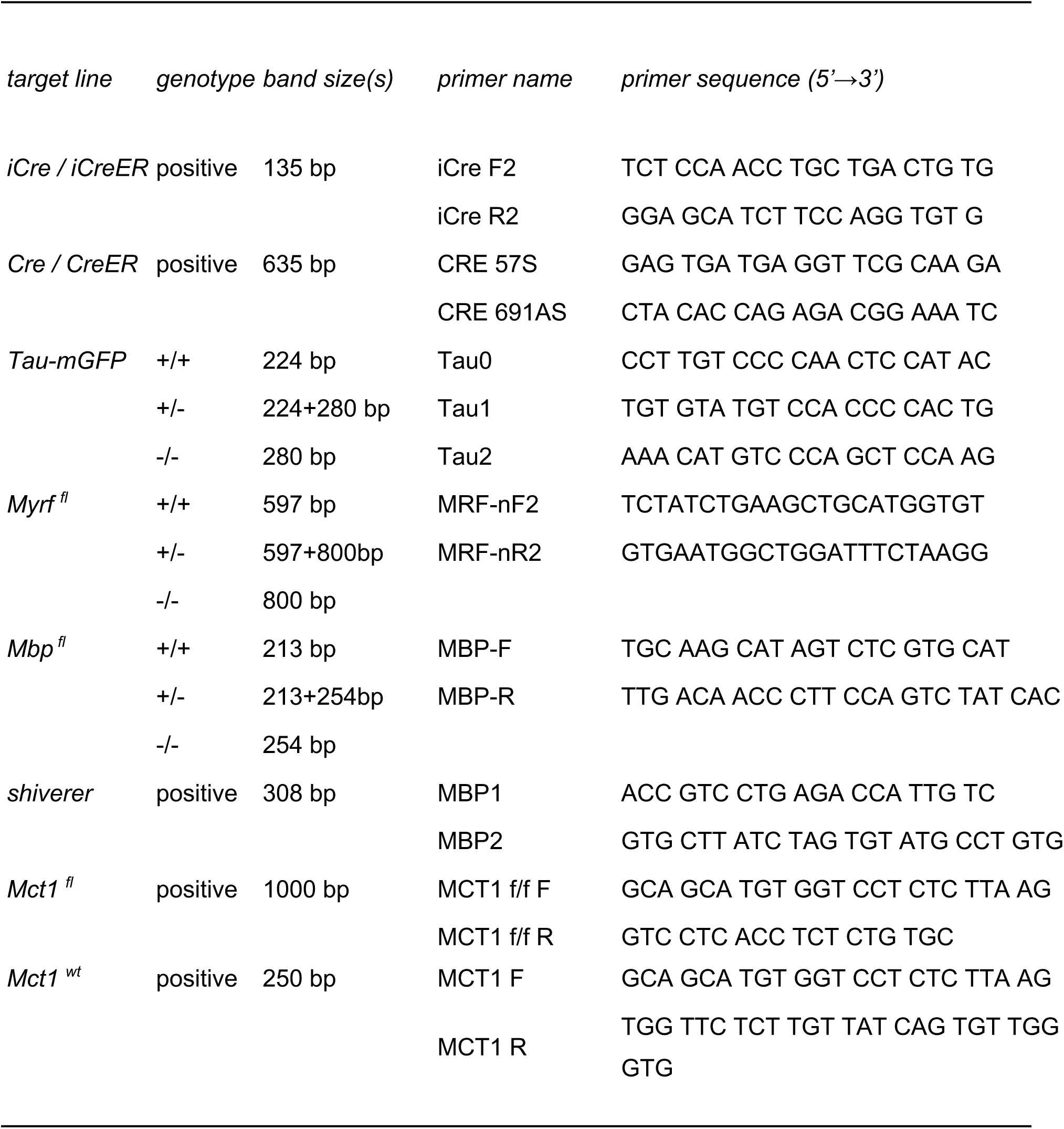
PCR primers for mouse genotyping.

**Supplementary Table S2.** Primary antibodies.

| *Antigen* | *Raised in* | *Dilution* | *Supplier* |
| --- | --- | --- | --- |
| MBP | rat | 1:300 | Abcam |
| CC1 | mouse | 1:200 | Millipore |
| CASPR1 | rabbit | 1:500 | Millipore |
| CASPR1 | mouse | 1:500 | Abcam |
| NaV1.6 | rabbit | 1:500 | Alamone Labs |
| pan-NaV | mouse | 1:500 | Sigma |
| GFP | chicken | 1:1000 | Aves Labs |

**Supplementary Movie S1:** Freely moving P28 *Sox10-Mbp-cKO* mouse with *Mbp^(fl/+)^* littermate.

**Supplementary Movie S2:** Freely moving P28 *Mbp^(shi/shi)^* mutant mouse.

**Figure S1.**
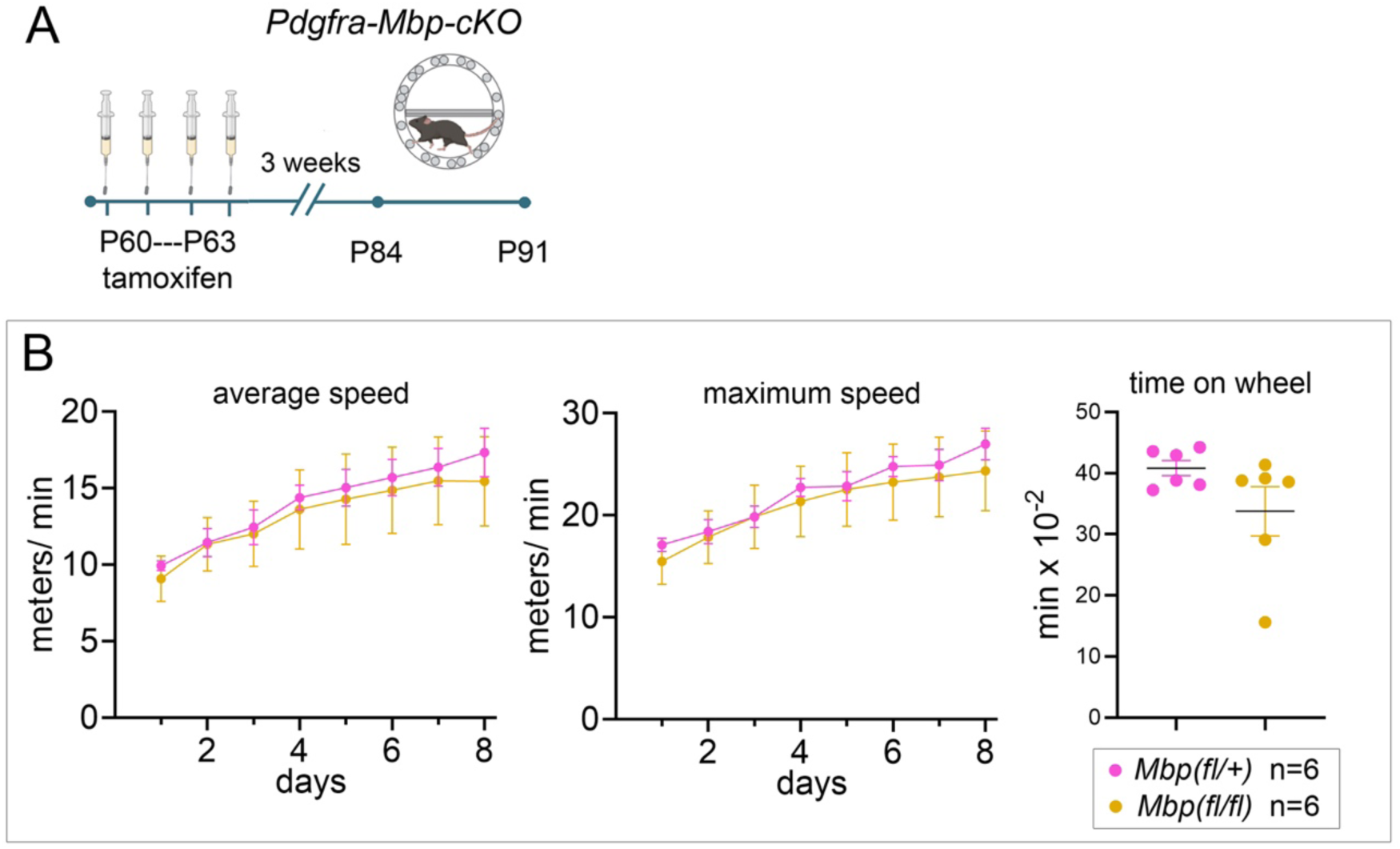
Intact motor skill learning in *Pdgfra-Mbp-cKO* mice. (A) Timeline of the experiment. Tamoxifen was administered to *Pdgfra-Mbp-cKO* mice and control littermates on P60–P63 and 3 weeks later they were introduced to the complex wheel for 8 days. (B) Running speed was recorded automatically during the active periods and daily average and maximum speeds were calculated. There were no significant differences in running performance between the two cohorts, either in average or maximum speed over the 8 days housed with the wheel. Time spent turning the wheel was reduced in *Pdgfra-Mbp-cKOs* relative to controls although this did not reach statistical significance.

**Supplementary Figure S2.**
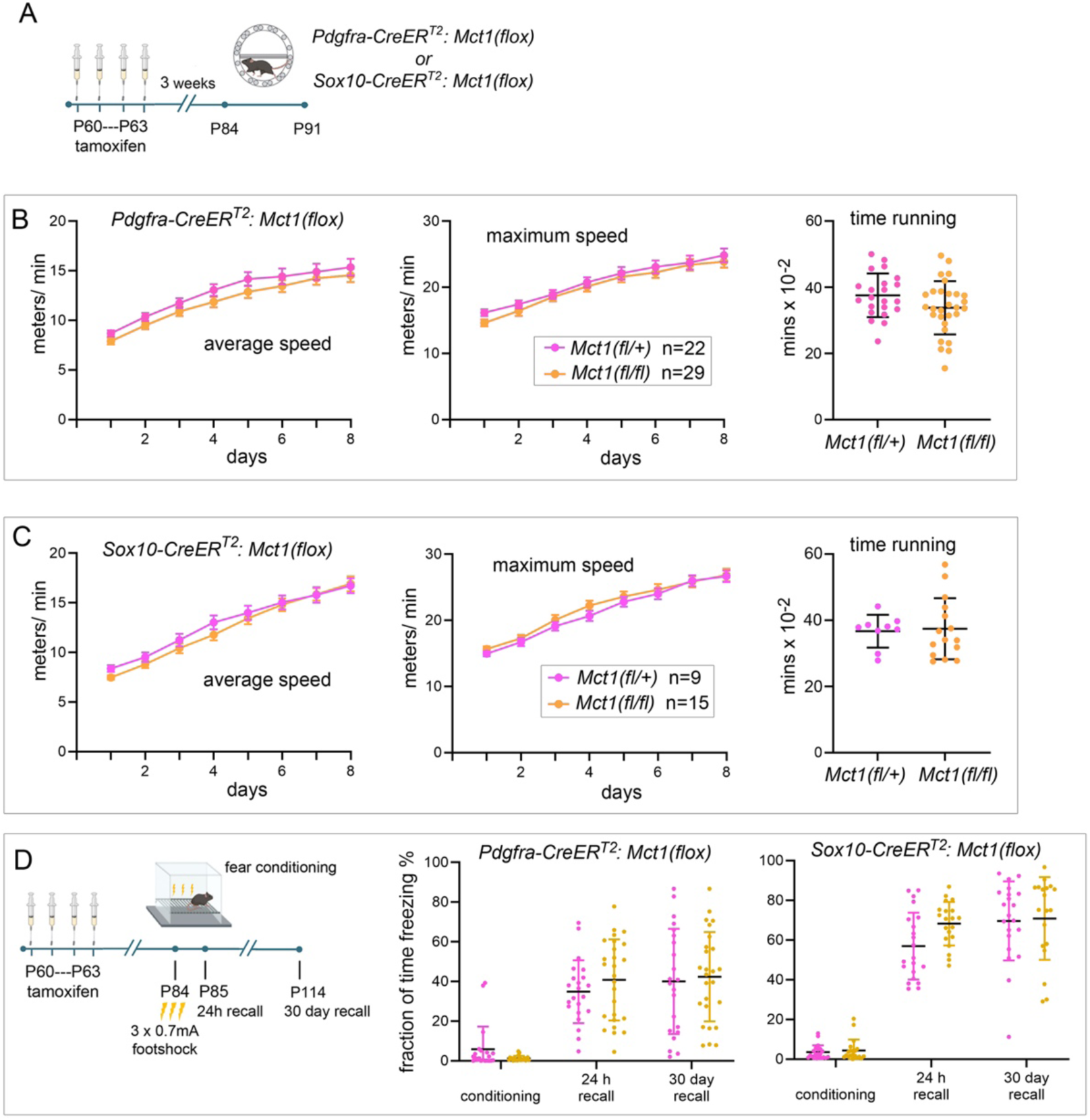
Intact learning and memory consolidation in *Mct1-cKO* mice. (**A**) Experimental timeline. Tamoxifen was administered to *Pdgfra-Mct1-cKO* or *S10-Mct1-cKO* mice and control littermates on P60–P63, then after a 3-week recovery period they were introduced to the complex running wheel for 8 days. (**B**, **C**) The running performances of both *Pdgfra-Mct1-cKO* (**B**) and *S10-Mct1-cKO* mice (**C**) were indistinguishable from controls. Both *cKO*s spent the same amount of time tuning the wheel as their control littermates. (**D**) *Pdgfra-Mct1-cKO* and *S10-Mct1-cKO* mice underwent contextual fear conditioning, along with control littermates. Their freezing behaviour was recorded during the conditioning period and again at 24 h and at 30 days after conditioning. No significant differences were observed between *Pdgfra-Mct1-cKO* or *S10-Mct1-cKO* mice and their control littermates.

